# Computational Electrostatics Predict Variations in SARS-CoV-2 Spike and Human ACE2 Interactions

**DOI:** 10.1101/2020.04.30.071175

**Authors:** Scott P. Morton, Joshua L. Phillips

## Abstract

SARS-CoV-2 is a novel virus that is presumed to have emerged from bats to crossover into humans in late 2019. As the global pandemic ensues, scientist are working to evaluate the virus and develop a vaccine to counteract the deadly disease that has impacted lives across the entire globe. We perform computational electrostatic simulations on multiple variants of SARS-CoV-2 spike protein s1 in complex with human angiotensin-converting enzyme 2 (ACE2) variants to examine differences in electrostatic interactions across the various complexes. Calculations are performed across the physiological pH range to also examine the impact of pH on these interactions. Two of six spike protein s1 variations having greater electric forces at pH levels consistent with nasal secretions and significant variations in force across all five variants of ACE2. Five out of six spike protein s1 variations have relatively consistent forces at pH levels of the lung, and one spike protein s1 variant that has low potential across a wide range of pH. These predictions indicate that variants of SARS-CoV-2 spike proteins and human ACE2 in certain combinations could potentially play a role in increased binding efficacy of SARS-CoV-2 *in vivo*.

## 1 INTRODUCTION

SARS-CoV-2 (*CoV-2*) is the latest novel coronavirus to have crossed the species boundary into humans over the past twenty years [7, 22]. Currently available information strongly suggests an originating source of the CoV-2 virus to be the bat coronavirus SL-CoV-RaTG13 with 96% similarity to CoV-2 [21].

The World Health Organization (*WHO*) declared CoV-2 a public health emergency on January 30, 2020 and then pandemic status in the weeks following. WHO utilizes a tool to distinguish diseases based on the potential to bloom into epidemic/pandemic proportions and the availability of countermeasures to focus research and development (*R&D*) resources accordingly [19, 20].

We have previously performed extensive research using electrostatic analysis of HIV gp120 envelope glycoprotein interactions with human CD4 and broadly neutralizing antibody proteins [5, 6, 11–13, 15]. Electrostatics are a novel way to evaluate protein interactions at the structural level with a focus on how environmental conditions, like pH, effect binding efficacy. The processes can easily be adapted to any other protein-protein interactions which may be strengthened or weakened by modulating pH. The theoretical approaches have been further extended to predict electric force (Coulombs Law) modulation by pH using the same set of data generated by the electrostatic surface charge (*ESSC*) pipeline [10, 15].

## 2 RESULTS

For this research we utilized CoV-2 spike (s1) proteins: 6LZG_B, 6M17_E, 6VW1_2, 6W41_C, QIS30425.1, QIS60489.1 and angiotensin--converting enzyme 2 (*ACE2*) proteins: AAQ89076.1, BAG37592.1, BAJ21180.1, EAW98891.1, XP_011543854.1 obtained through sequence similarity searches using NCBI Protein BLAST and the Protein Data Bank [2]. Structurally, CoV-2 s1 shares similar traits with SARS-CoV s1 that emerged in 2003 with a major point of difference being an additional cleavage in the spike s1 protein sub-unit [17] making the substitution of similar SARS-CoV species difficult to justify in modeling and simulation work.

Computational methods of the pipeline performed as expected on s1 (see methods), but some results differed in content greatly when compared to those of HIV gp120. Past investigations predict stronger binding affinity between HIV gp120 and human CD4 proteins at lower physiological pH compared to higher physiological pH. Evidence of this was demonstrated by the pipeline EFP and BE data from those protein and complexes [5, 6, 11, 13]. However, the EFP and BE data for s1-ACE2 interactions predict no notable differences in binding efficacy across the physiologically relevant pH range. These data suggest that the s1-ACE2 interaction is therefore not pH-sensitive. For example, in Figure 1 we present a typical EFP for s1 and observe no notable shifts (away form zero) for these data in the physiologically relevant pH range. Color bars indicate the pH range of normal nasal secretions (red), pH 5.5 to 6.5; yellow indicates inflamed nasal secretions (rhinitis) from pH 7.2 to 8.3 [4]. Blue indicates the approximate lung pH listed as 7.38 to 7.43 [3], however, the graphs scale in 0.1 increments and displays the range as pH 7.3 to 7.5 as inclusive of the specific range. In Figure 2 we present the BEs for complexes showing the least and most variation, again observing no notable shifts for these data in the same relevant pH range.

**Figure 1:**
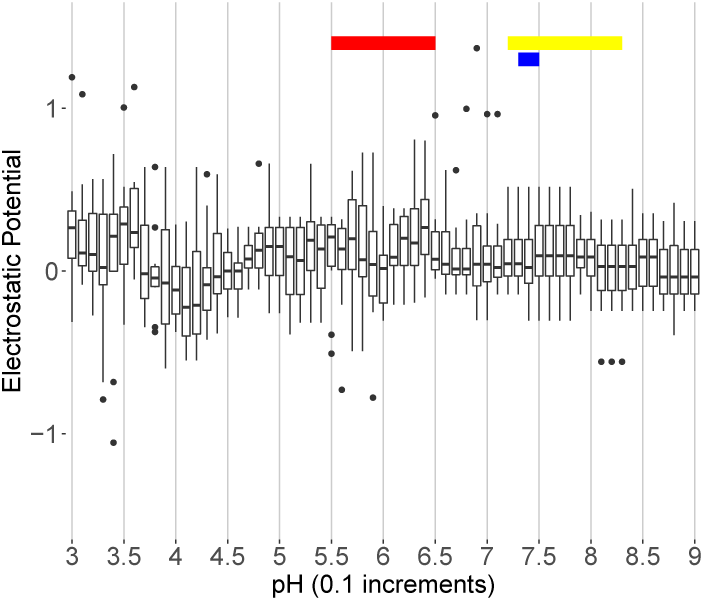
Typical electrostatic fingerprint of s1 subunit of CoV-2 envelope spike. Color bars near the top indicate the pH range of human nasal secretions (red), inflamed nasal secretions (yellow) and lung pH (blue). the fingerprint is basically flat from pH 5.0 and up indicating a stable structure. Color bars indicate pH range of normal nasal secretions (red), inflamed nasal secretions (yellow), and lung pH (blue).

**Figure 2:**
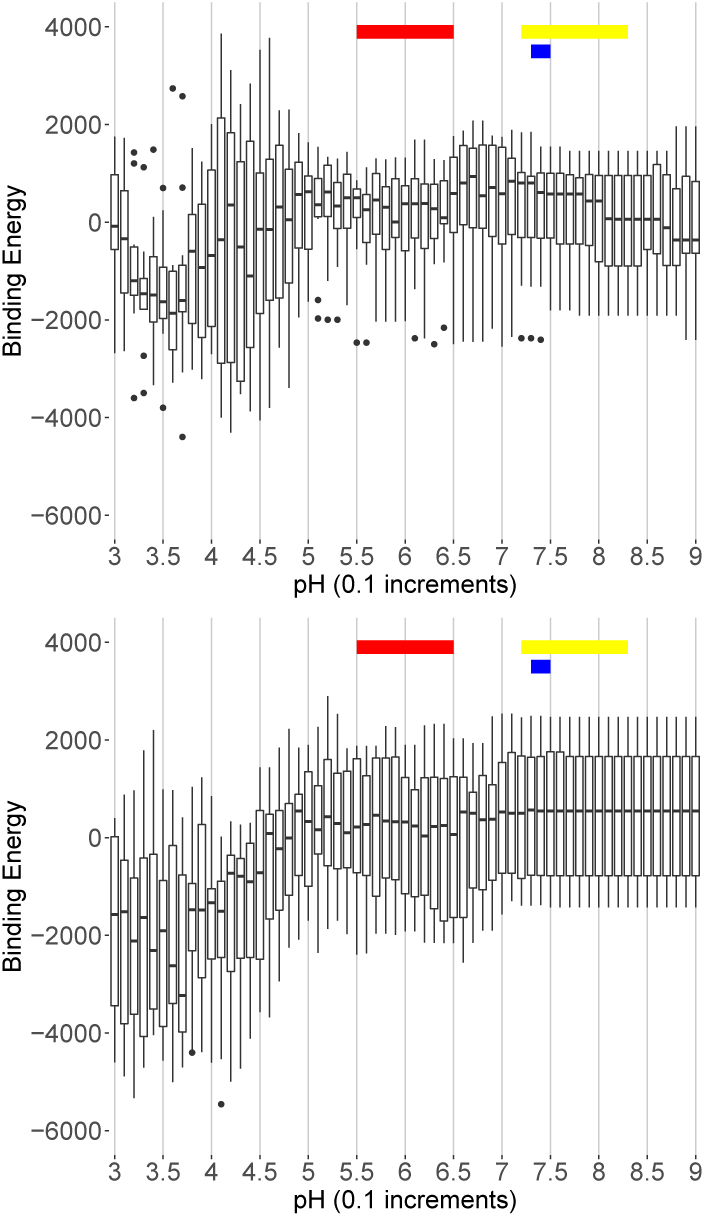
Predicted electrostatic contribution to binding energy across the physiological pH range shows some activity around nasal and lung pH levels in the most active plot (top), while a nearly flat result across the interesting range of pH 5.0 to 8.5 is observed in the most inactive plot (bottom). Color bars indicate pH range of normal nasal secretions (red), inflamed nasal secretions (yellow), and lung pH (blue).

One possible explanation for this lack of pH sensitivity is the minimal conformational change exhibited by s1-ACE2 interactions compared to gp120-CD4 interactions. The gp120-CD4 interaction requires a relatively large conformational change of the gp120 protein, leading to changes in the protein surface (residues) exposed to the external environment. In particular, an increase in negative surface change in the bound conformation (relative to the unbound conformation) of gp120, is thought to make the bound conformation energetically more favorable at low pH. However, the conformational change observed in s1-ACE2 complex binding appears to be either too small to alter, or simply results in too little alteration in the electrostatic surface charge of s1.

EFP and BE results predict that no or minimal differences exist in binding efficacy across the physiological pH range for s1-ACE2 interactions. These predictions leave unanswered questions, for instance, does the lack of differences in binding efficacy across the physiological pH range mean that s1 binds consistently regardless of pH? Another question, more pertinent to this research, what measurable forces are present between s1 and ACE2 that drive an interaction to taker place?

As a solution to the latter question, we expand the analysis with an alternative approach to electrostatic fingerprinting by taking the absolute values of the difference between s1 and ACE2 unbound conformation ESSC pipeline data to expose a predicted electric potential difference. We present results containing the highest and lowest potential differences from pH 4.5 to 9.0 in Figure 3. These data show an electric potential difference which varies across the examined pH range exists of varying quantity and across all complex combinations evaluated. Furthermore, the pipeline data predicts that s1 is positively charged throughout the pH range while ACE2 has a transition to negative between pH 4.5 and 5.0 as show in Figure 4. The presence of electric potential differences allow the results to be interpreted in terms of electric force.

**Figure 3:**
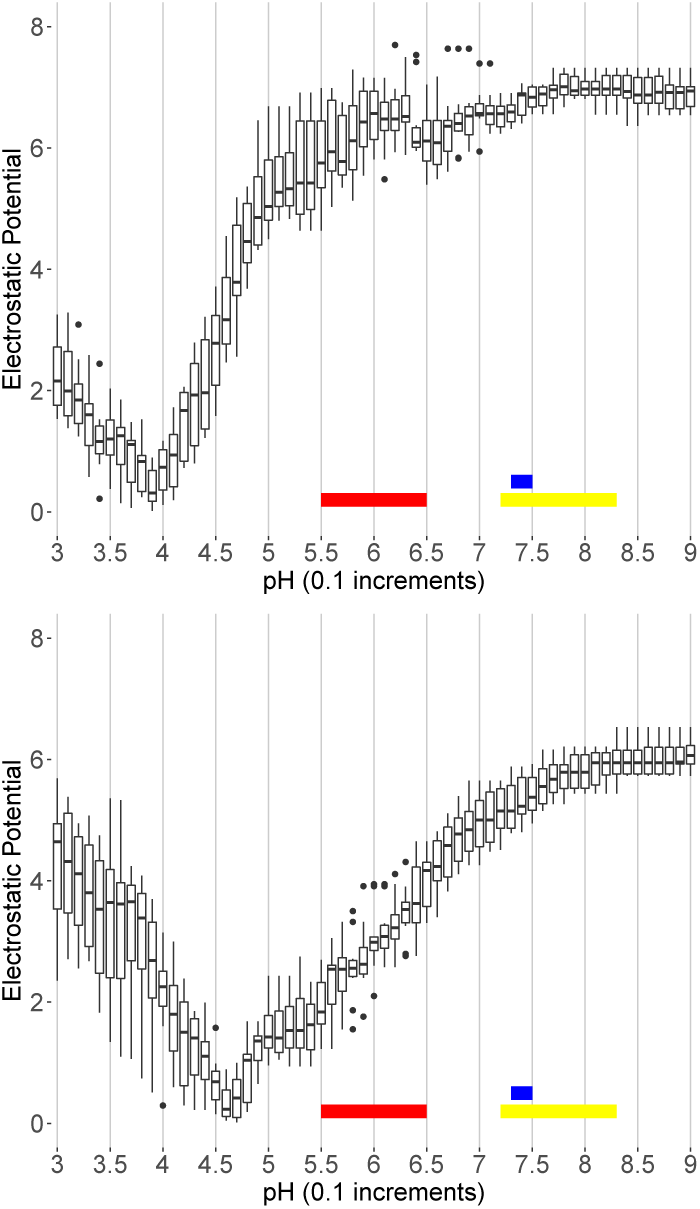
Comparison of potential differences between s1 and ACE2 at result extremes contrasting the highest potential differences (top) with the lowest (bottom). Color bars indicate pH range of normal nasal secretions (red), inflamed nasal secretions (yellow), and lung pH (blue).

**Figure 4:**
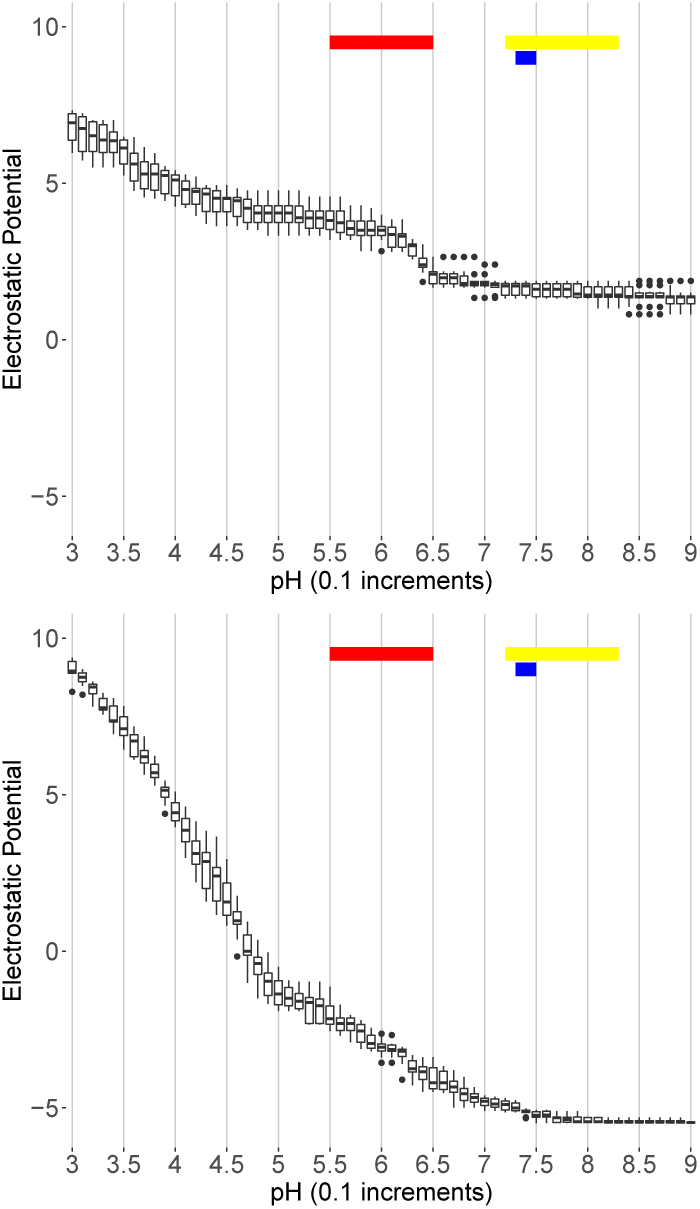
Typical ESSC pipeline results for s1 (top) and ACE2 (bottom). In all cases, s1 never crosses zero while ACE2 always transitions into negative values. Color bars indicate pH range of normal nasal secretions (red), inflamed nasal secretions (yellow), and lung pH (blue).

The presence of electric potential difference introduces a force that can be determined by applying Coulombs Law to obtain the Electric Force (*F*_*e*_) in Newtons (*N*). We can graph the changing forces, either repulsive (+) or attractive (-) between s1 and ACE2, across pH range of 3.0 to 9.0 in 0.1 increments at a distance of 10Å. Our model is *non-specific* in that specific interactions involve close-proximity contacts between proteins such as hydrogen bonds, unlike charges, and hydrophobic residue stacking. However, non-specific interactions are not close-proximity, and can be attractive or repulsive beyond the range of 3 angstroms or more. Electric force is a good example of the latter case. Details of the calculation are provided in the Methods section below.

Figure 5 displays the electric force from pH 3.0 to 9.0 in 0.1 increments for all complexes. Positive values indicate a repulsive force and negative values indicate an attractive force. For Figure 6 we discard the repulsive forces for the sake of figure clarity and on the basis that they exist outside the relevant pH range.

**Figure 5:**
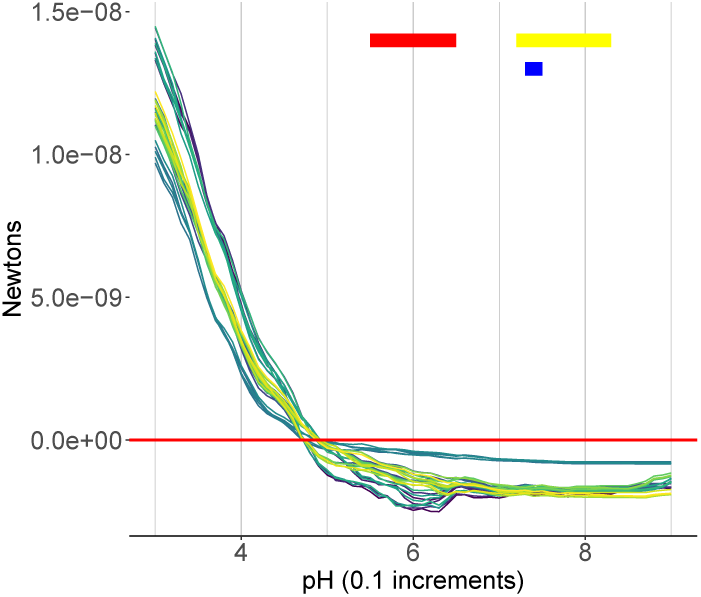
Aggregation of electric force for all complexes. Positive values are repulsive and negative values are attractive. Color bars indicate pH range of normal nasal secretions (red), inflamed nasal secretions (yellow), and lung pH (blue).

**Figure 6:**
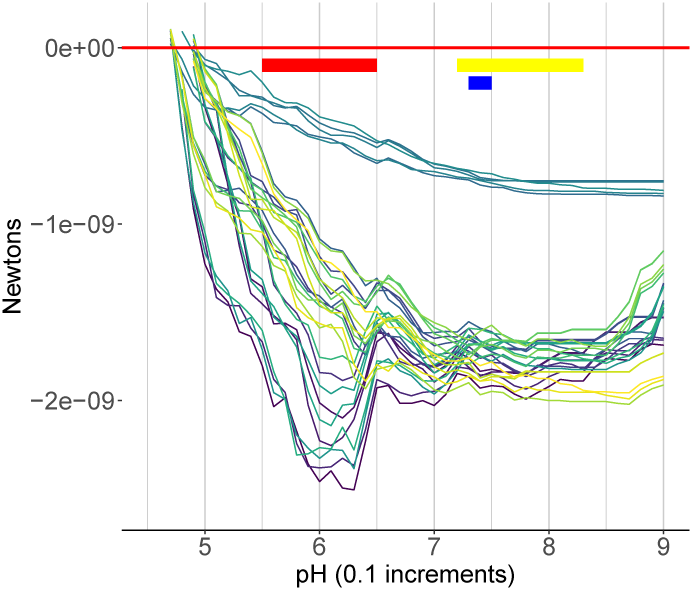
Aggregation of electric force for all complexes analyzed minus repulsive forces. This representation allows for a broad interpretation of the data where a single group of complexes stands out from the rest and another group has large spikes at nasal secretion pH levels. Color bars indicate pH range of normal nasal secretions (red), inflamed nasal secretions (yellow), and lung pH (blue).

In Figure 6 we can see unique characteristics emerging where a group of complexes stand out from the rest with weakly attractive forces in a smooth curve across the examined range; another cluster of complexes emerges with a higher level of force in a slightly more erratic curve across the range and a group of complexes with a pronounced peak in force across the range of nasal secretions that peaks in the approximate middle.

While more details can be obtained from Figure 6, a clearer understanding is provided by Figure 7, which is grouped by s1. Clear patterns emerge that indicate higher forces presents for s1 proteins 6LZG_B and 6W41_C at nasal secretion pH. Additionally, ACE2 genetic variations AAQ89076.1 and EAW98891.1 are most impacted by these two s1 variants. However, 6M17_E, QIS30425.1, and QIS60489.1 have nominal forces across the interesting pH range with marginal differences among ACE2 variants. 6VW1_2 has low potential across the pH range and little differences across ACE2 variants.

**Figure 7:**
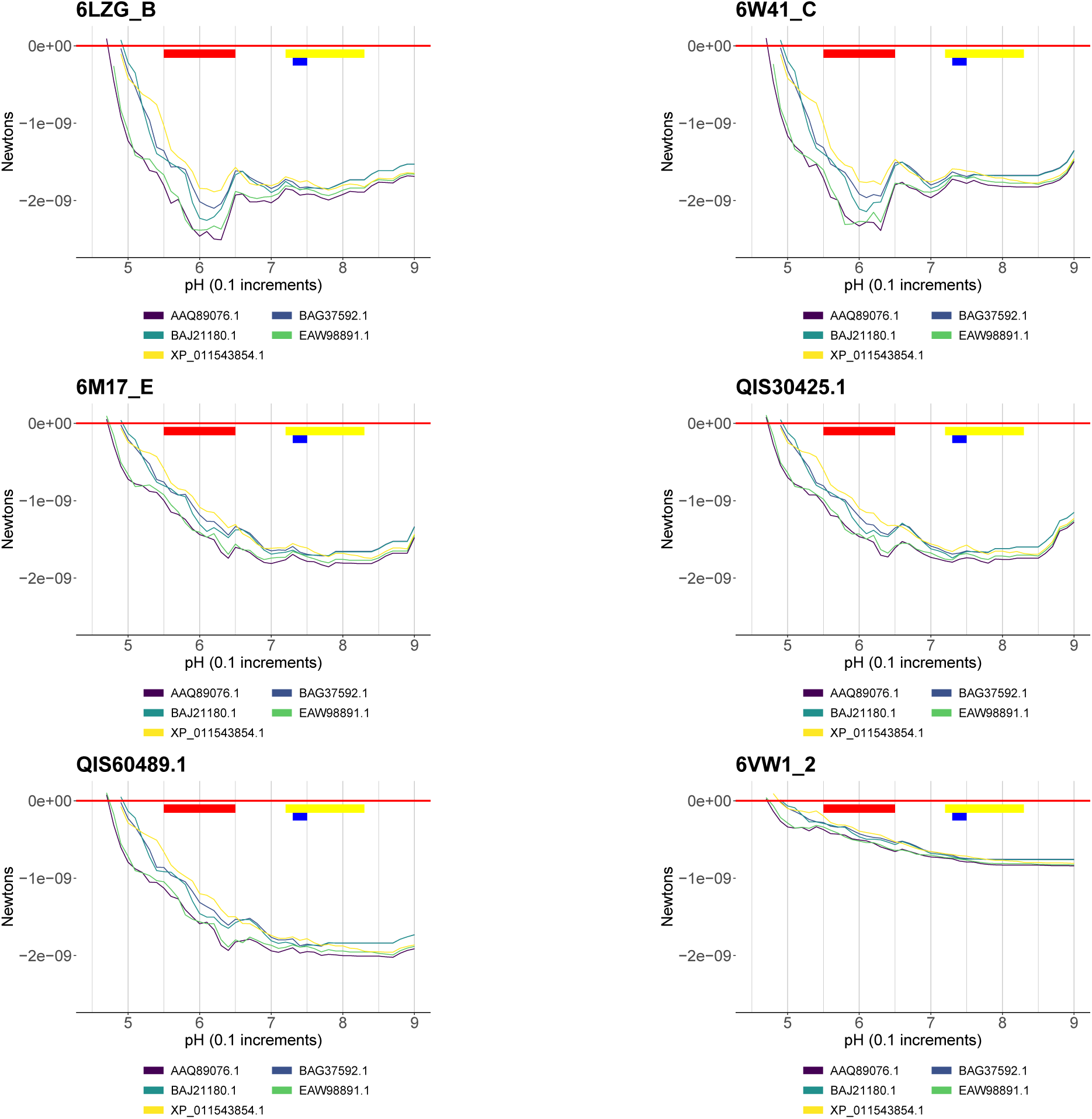
Grouped by s1, patterns emerge that indicate higher forces for s1 proteins 6LZG_B and 6W41_C at nasal secretion pH and distinct bias for ACE2 variants AAQ89076.1 and EAW98891.1, a lesser bias for ACE2 BAJ21180.1 and BAG37592.1. CoV-2 variants 6M17_E, QIS30425.1, and QIS60489.1 have nominal forces across the interesting pH range with marginal differences among ACE2 variants. 6VW1_2 has lower forces across the pH range and little differences across ACE2. Color bars indicate pH range of normal nasal secretions (red), inflamed nasal secretions (yellow), and lung pH (blue).

## 3 DISCUSSION

Our results clearly predict that sequence variations of s1 and ACE2 directly impact the attractive forces involved between s1 and ACE2. Furthermore, these results implicate human ACE2 variations that exacerbate those forces for specific s1 and ACE2 combinations. Additionally, we point out that three groupings emerge in this analysis showing two s1 variants (6LZG_B and 6W41_C) with large spikes in attractive force at pH 5.5 to 6.5 associated with nasal secretions. Three s1 variants (6M17_E, QIS30425.1, and QIS60489.1) have a relatively smooth curve at nominal attractive forces across ACE2 variants and one s1 variation (6VW1_2) has a smooth curve with low attractive force and minimal differences across ACE2 variations.

Not all s1 and ACE2 protein sequences examined have solved crystal structures and were instead modeled based on the currently available s1 and ACE2 structures (see methods). Additional crystal structures for either or both s1 and ACE2 would allow for more precise predictions. Additionally, an analysis of electric force at the amino acid level would provide more detailed predictions at specific regions involved with s1 and ACE2 contact points and will be the subject of future studies.

We provide access to the full set of figures and results generated from this study as a tar package located at: https://github.com/spmorton/SARS-CoV-2

## 4 METHODS

All computational methods related to the electrostatic pipeline are detailed in [5, 6, 11–13, 15]. Templates used to model s1 are: 6LZG, 6M0J, 6VW1, and 6VYB [9, 14, 17, 18] and for ACE2 the templates are: 1R4L, 1R42, 6M0J, 6VW1 [9, 14, 16]. To target Frodan for specific conformations of s1 we used: 6LGZ as bound and 6VYB as unbound target conformations. For ACE2 we used 6LZG for bound and unbound target conformations.

For the specific calculation of Electric Force, APBS returns a factor for *V* in:

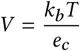

Where *V* is voltage of the system per unit, *k*_*b*_ is the Boltzmann’s constant: 1.3806504 E-23 *J K* ^−^1, *T* is the temperature in Kelvin and *e*_*c*_ is the charge of an electron: 1.60217646 E-19 C.

For the calculation of Electric Force (*F*_*e*_) we have:

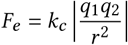

Where *F*_*e*_ is the Electric Force in Newtons, *k*_*c*_ is Coulombs constant: 8.987551E+09, *q*_1_ is the charge in Coulombs of the first mass, *q*_2_ is the charge in Coulombs of the second mass, and *r* is the distance in meters between the two masses.

To derive Coulombs of charge from APBS [1, 8] requires a simple transposition of the terms *V* and *e*_*c*_ and multiplication of the results with values returned by APBS. For this model, the variables required to complete the calculations are: *T* = 310*K* and *r* = 10Å.

